# Maintenance of polymorphism in spatially heterogeneous environments

**DOI:** 10.1101/2025.04.25.650679

**Authors:** Takahiro Sakamoto, Sam Yeaman

**Affiliations:** Department of Biological Sciences, University of Calgary, Calgary, AB T2N 1N4, Canada; National Institute of Genetics 1111 Yata, Mishima, Shizuoka 411-8540, Japan

**Keywords:** population genetics, local adaptation, spatial model, theory

## Abstract

Local adaptation occurs when species adapt to spatially heterogeneous environments. The stability of local adaptation is determined by selection-migration-drift balance: selection favours adaptive divergence whereas migration and random genetic drift cause the collapse of the divergence. How this balance determines the dynamics of local adaptation has been extensively studied. However, owing to the complexity of multi-population models, most previous theories used models with simple population structure and environmental variation, limiting their applicability to complex situations in nature. To address this issue, we developed a new theoretical method to analyze complex multi-population models, allowing heterogeneity in selection, migration, and population density. In essence, our method approximates a spatial model by a panmictic one-population model while retaining the core stochastic structure, enabling the application of conventional diffusion methods. By comparing with simulations, we confirmed that our method accurately describes stochastic evolutionary dynamics in various spatial models, provided sufficient levels of migration. This method is then applied to examine the effect of environmental distribution in 2D space. Assuming landscapes with different levels of the spatial autocorrelation of the environment, we found that the maintenance of locally adaptive alleles is significantly promoted when the spatial autocorrelation is high. These results highlight how complex spatial heterogeneity, as seen in nature, could affect the qualitative outcome of evolution.

---

Spatially varying selection is a major mechanism for maintaining genetic diversity. Under strong divergent selection, different alleles dominate in different environments, coexisting at migration-selection balance. In finite-size populations, however, random genetic drift causes stochastic fluctuation of allele frequencies, potentially leading to the loss of locally adaptive alleles. Constructing a theory to predict the persistence of adaptive divergence is crucial not only for our fundamental understanding of genetic variation, but also for practical issues such as conservation of intraspecific variation (Des Roches *et al*. 2018, Des Roches *et al*. 2021; Meek *et al*. 2023) and prevention of unfavored local adaptation like evolution of drug resistance (Kliot and Ghanim 2012; Hawkins *et al*. 2019).

Natural environments are inherently complex in their patterns of spatial variation, but previous theories do not fully capture this complexity. Where deterministic models have been used to examine the effect of environmental heterogeneity (Haldane 1930, 1948; Fisher 1950; Deakin 1966; Slatkin 1973; Nagylaki 1975; Akerman and Bürger 2014), they have ignored random genetic drift and therefore do not capture important consequences of stochasticity inherent in real populations. On the other hand, previous stochastic models have often focused on simple patterns of environmental variation, such as two-patch (Wright 1931; Yeaman and Otto 2011; Aeschbacher and Bürger 2014; Yeaman *et al*. 2016; Sakamoto and Innan 2019) and island models (Pollak 1966; Barton and Whitlock 1997; Szép *et al*. 2021); thus, these models do not represent the true complexity and non-linearity of real environments. Critically, even for relatively simple scenarios, the persistence of locally adaptive divergence remains elusive due to the complexity of the multi-population model. Most studies have focused on establishment of a new locally adaptive allele (Pollak 1966; Barton 1987; Yeaman and Otto 2011) and did not consider the stochastic dynamics after the evolution of local adaptation. So far, the persistence is well quantified only for the continent-island model (i.e., two patches with unidirectional migration) (Aeschbacher and Bürger 2014).

In this study, we aim at evaluating the stability of locally adaptive divergence allowing more realistic spatial heterogeneity. To this end, we develop a novel method to approximate the evolution in a multi-population model using a one-dimensional diffusion process. Our theoretical idea is illustrated in Figure 1A, where, as an example, allele frequencies observed in a two-population model under divergent selection are plotted. In the simulation, we first randomly assigned a mutant allele to one of the two populations and ran simulations many times to obtain frequency trajectories. Allele frequencies in each population can diverge significantly due to limited migration and divergent selection, but they do not evolve independently and are consistently trapped around the frequency path of the corresponding deterministic model (Figure 1A). The existence of this typical trajectory inspires an idea that the evolutionary process can be approximated by a one-dimensional diffusion on the deterministic path.

**Figure 1.**
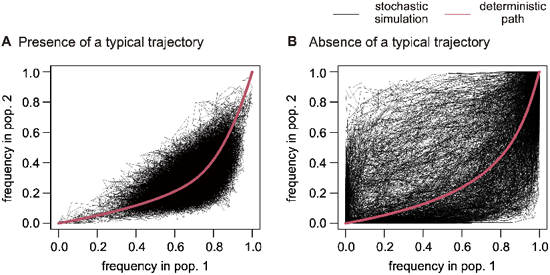
Frequency trajectories in the stochastic simulations of two-subpopulations model. Parameters are 2*N* = 250, *s*_1_ = − 0.04, *s*_2_ = 0.04, and *m* = 0.02 in (A), and 2*N* = 250, *s*_1_ = 0.05, *s*_2_ = 0.03, and *m* = 0.005 in (B).

Unfortunately, in some cases, the evolutionary trajectories do not follow a typical trajectory. An example of this is shown in Figure 1B, where small *m* and globally positive selection are assumed. In this case, the focal allele is likely to increase rapidly in one of the populations (usually in the population where it originates), thus no typical frequency trajectory exists. In other words, the evolutionary process is intrinsically multidimensional and cannot be reduced to the one-dimensional process. This pattern suggests that our approximation requires that migration is high enough that allele frequencies in each population are well-coupled. One may consider this condition restrictive, but below we show that intermediate migration rate is often sufficient and this strategy works in many scenarios with local adaptation.

In the following, we first present theoretical methods for judging whether a typical frequency trajectory exists so that the method is applicable, and calculating diffusion coefficients for applicable cases. The predictions based on the constructed diffusion processes are compared with the simulation results, showing a general utility of the present method. We then apply our method to explore how different spatial distribution of environments affect the stability of local adaptation, and highlight that the spatial autocorrelation of environment greatly improves the stability of divergence. Although this pattern has been qualitatively known from deterministic theory and simulations for some scenarios (Slatkin 1973; Svoboda *et al*. 2023; Booker 2024), our theory is a first study to provide a mathematical formulation for quantitative predictions on stochastic dynamics which is applicable to a wide range of patterns of heterogeneity in environment, migration, and population density.

It should be noted that this method is different from previous perturbation-based approaches (Whitlock and Gomulkiewicz 2005; Constable and McKane 2014) where selection is considered to induce a small frequency deviation from neutral processes. Although the perturbation approaches allow small heterogeneity of selection as long as the frequency divergence among demes is subtle, they cannot be applied to cases with large frequency divergence, such as local adaptation (see SI Section 3 for the detailed discussion). Our theoretical approach aims at analyzing cases with large allelic divergence including local adaptation, for which no analytical tools were previously available.

## Results

### Model

We consider a one-locus *n*-subpopulation model. 2*N*_*i*_ haploid (or *N*_*i*_ diploid) individuals exist in *i*th subpopulation. There are two alleles (allele A and a) at the locus, where log fitness of alleles A and a in *i*th subpopulation is denoted by *s*_*i*_ and 0, respectively. For diploid species, codominance (additivity) is assumed. Backward migration rate from *j*th subpopulation to *i*th subpopulation (i.e., proportion of individuals in subpopulation *i* that migrated from subpopulation *j*) is *m*_*ij*_. The frequency of allele A in *i*th subpopulation is denoted by *p*_*i*_.

### Diffusion construction

The procedure to construct a diffusion process is illustrated in Figure 2 (see SI Section 1 for details). First, assuming that the typical frequency trajectory can be approximated by the deterministic frequency path, we calculate it as a function of average allele frequency, 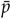 (Figure 2A). In this example, 2*N*_1_ = 2*N*_2_ = 1,000, *s*_1_ = 0.02, *s*_2_ = − 0.02, *m*_12_ = *m*_21_ = 0.01 is assumed. Next, we check whether evolutionary trajectories starting nearby the deterministic path tend to approach the path for each 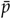 (Figure 2B). To quantify this, we project the deterministic evolutionary direction (i.e., 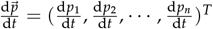) onto the plane that is orthogonal to the deterministic path and check if the projected vector is toward the deterministic path (*λ* < 0) or not (*λ ≥* 0). When *λ* < 0 for all 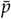, we consider that the evolutionary dynamics are trapped around the deterministic path and reducible to the one-dimensional process. For those cases, we calculate diffusion coefficients 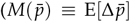 and 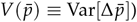 for each 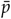 (Figure 2C) using an approximation based on the evolutionary dynamics nearby the deterministic path. This diffusion method is expected to work when evolution starts from a frequency set that is close to the deterministic path. However, it may not work for new mutations because they typically exist in one of the subpopulations and do not have a geographically balanced distribution. To address this issue, we utilized the branching process approximation to determine “effective” initial frequency 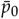 for new mutations. In practice, we determine 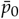 such that our constructed diffusion gives the same establishment probability as the one derived by the branching process (Pollak 1966; Barton 1987). Throughout the paper, for new mutations, we restrict our focus on mutations with positive invasion fitness because only those mutations should avoid immediate extinction and show non-trivial dynamics. These values 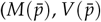, and 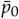) are substituted into the one-dimensional diffusion theory (Crow and Kimura 1970; Ewens 2004) to predict the evolutionary dynamics.

**Figure 2.**
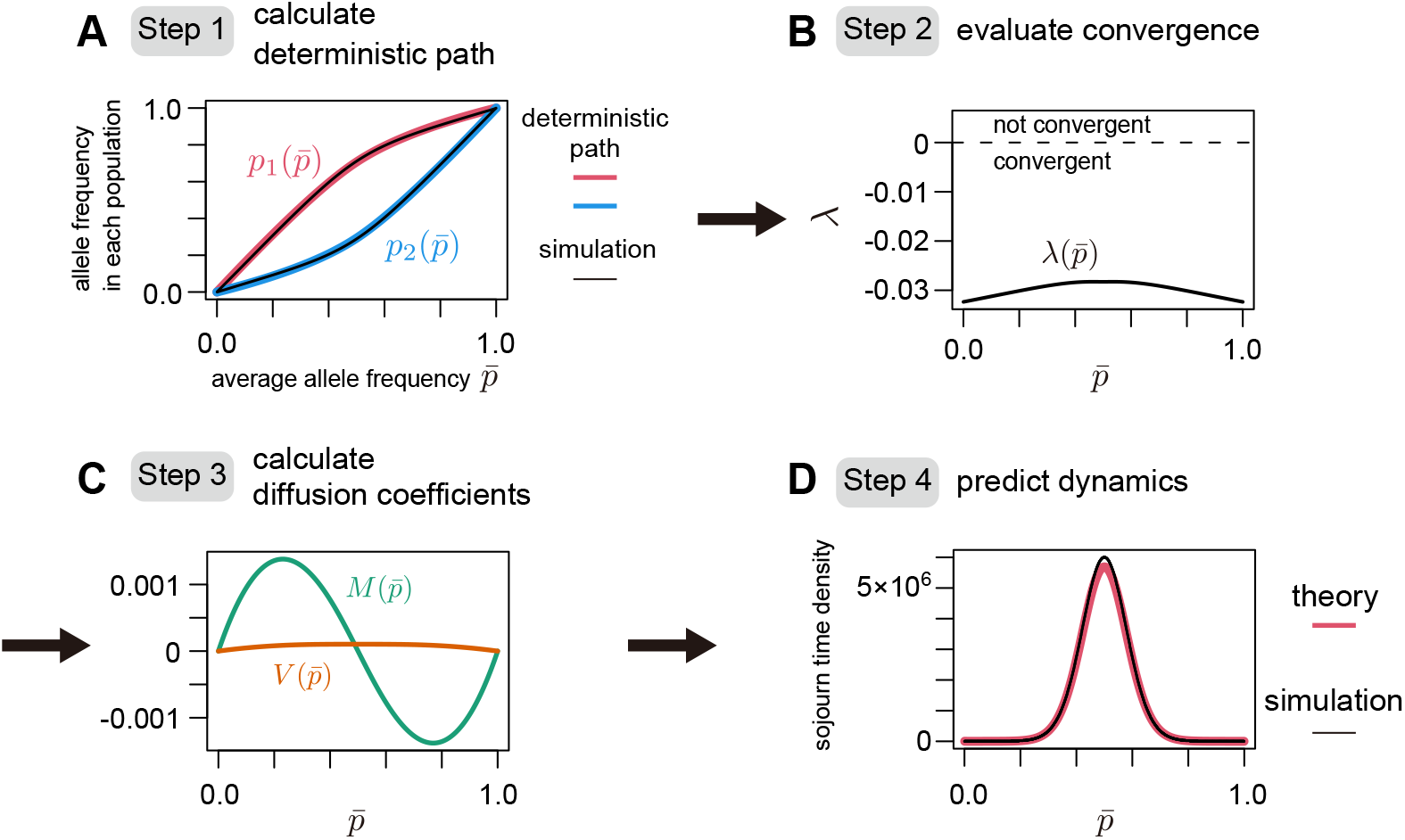
Illustration of our approximation method. In this figure, it is assumed that 2*N*_1_ = 2*N*_2_ = 1,000, *s*_1_ = 0.02, *s*_2_ = − 0.02, *m*_12_ = *m*_21_ = 0.01 as an example. (A) Firstly, we calculated the deterministic frequency path (red and blue lines). The average frequency in the simulations 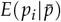 is also plotted (black line) for comparison. (B) Check whether evolutionary direction around the deterministic path directs to the path (*λ* < 0) or not (*λ*≥ 0) for each 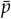. (C) Calculate the strength of deterministic and stochastic forces as functions of 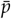. (D) Using obtained diffusion coefficients, the evolutionary dynamics are calculated. In this panel, sojourn time density is compared between the theory and simulation.

To showcase the validity of our approach, we ran stochastic simulations for the comparison in Figure 2, starting from *p*_1_ = 1/(2*N*_1_) and *p*_2_ = 0. Figure 2A shows that the deterministic path is a good approximation for the average trajectory observed in the stochastic simulations (Figure 2A). We also compare a sojourn time density at each frequency between the theory and simulation in Figure 2D. Theoretical values are calculated following diffusion theory for the one-population model (Crow and Kimura 1970; Ewens 2004) while 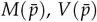, and 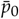 are used (see SI Section 2 for details). Figure 2D shows that the simulation results agree well with the theoretical prediction across the entire frequency range, suggesting that our approximation method accurately summarizes the core stochastic dynamics while simplifying the mathematical structure.

### Comparison between theory and simulation

To check the performance across wider parameter space, we calculated the mean absorption time of a new mutation (i.e., average time until either of the alleles is fixed in the entire population) in the two-population model (see SI Section 2 for derivation). The theoretical prediction and simulation results are compared while modifying *s*_1_ and *s*_2_ for three migration rates (Figure 3A– C). An initial state of *p*_1_ = 1/(2*N*_1_) and *p*_2_ = 0 was assumed. In each panel, color of the background squares represents the theoretical prediction for mean absorption time, while color of the inside circles shows the simulation results. The contour plot shows the convergence strength toward the deterministic path *λ*_max_ so that the theoretical result is unavailable when *λ*_max_ > 0 (colored in gray) due to the lack of convergence for some 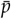. The simulation results are calculated as an average over 100,000 replicates. We stopped simulation runs when the number of generations exceeds 10^9^ and show parameters with those runs in white circles.

**Figure 3.**
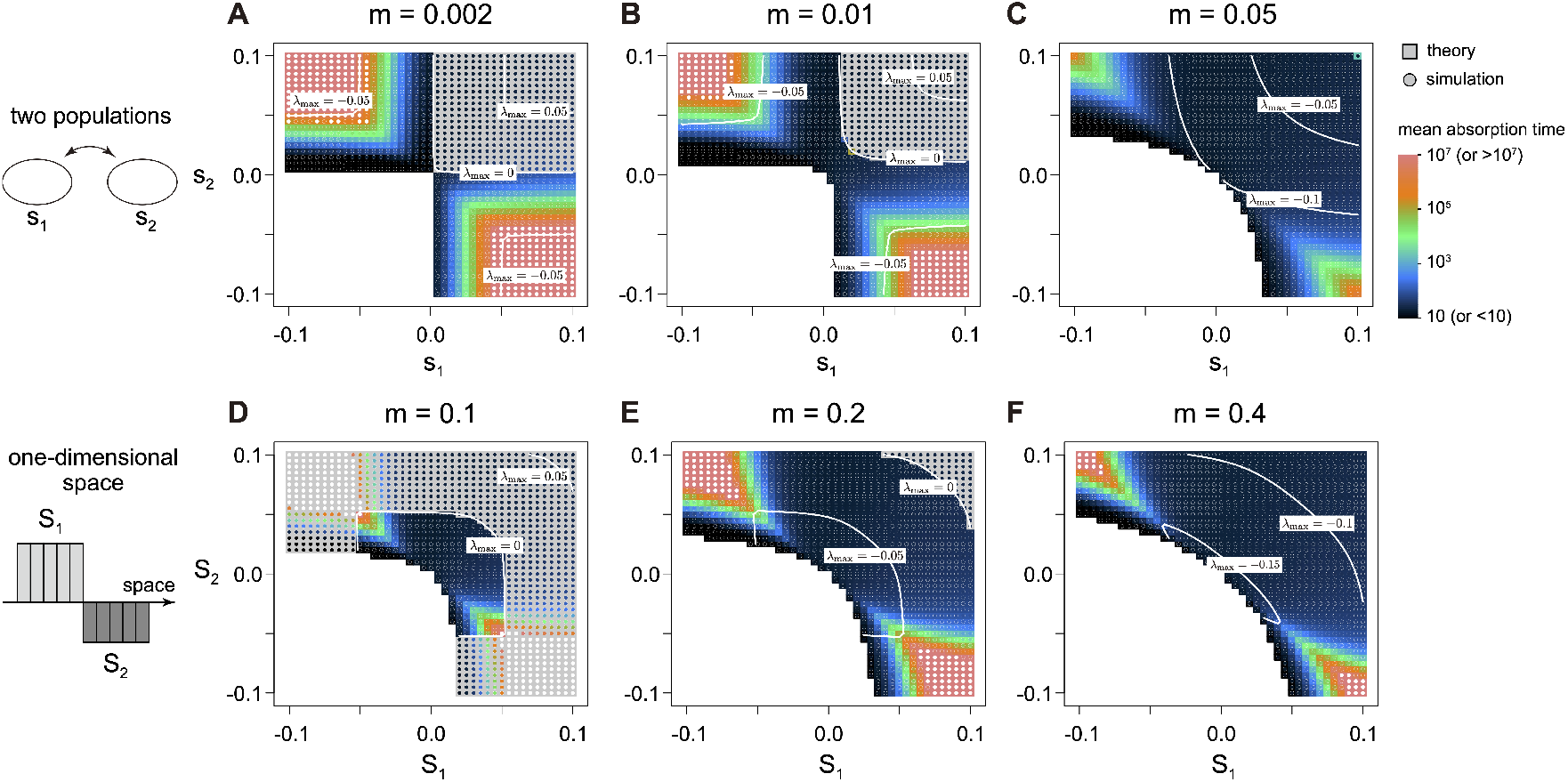
Mean absorption time in the two-population model (A–C) and one-dimensional spatial model (D–F). In (A–C), 2*N*_*i*_ = 200 was assumed, while in (D–F), 2*N*_*i*_ = 100 was assumed. White circles for simulation results mean that fixation does not occur within 10^9^ generations in some replicates.

Figure 3A–C shows how the absorption time depends on the selection strength. As expected, the absorption time becomes very long when *s*_1_ and *s*_2_ have opposite sign and *m* is small because divergent selection maintains adaptive divergence for a long time. The agreement between the square and circle colors shows that our theory accurately predicts the stochastic dynamics for applicable cases. We found that the parameter space to which the method is inapplicable (*λ*_max_ > 0, gray areas) expands as the migration rate decreases. This is because evolutionary processes in different subpopulations are more likely to be decoupled for small *m*. Inapplicability tends to arise when globally positive selection works, consistent with the observation in Figure 1B. In these cases, the focal allele can increase locally in either of the subpopulations, which explains the absence of typical trajectory. In contrast, our method generally works when divergent selection acts even at low migration rates (Figure 3A), suggesting its utility in cases with local adaptation. We checked our method’s performance assuming further heterogeneity in subpopulation size and migration asymmetry (Figure S3, S4), and confirmed the validity as long as *λ*_max_ is small enough (typically *λ*_max_ < − 0.01). We also compared the performance of our method with previous methods that are based on perturbation from neutral processes (Whitlock and Gomulkiewicz 2005; Constable and McKane 2014), and showed that our method outperforms in most parameters, especially for the local adaptation cases (SI Section 3).

We next extend our focus to more complex multi-deme models. We consider a one-dimensional space consisting of 10 demes, each of which belongs to one of the two environments (*E*_1_ or *E*_2_). Selection coefficient for allele A is denoted by *S*_1_ and *S*_2_ in *E*_1_ and *E*_2_, respectively. Migration rate between demes is assumed to be

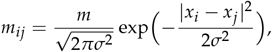

where *x*_*i*_ is the position of *i*th deme (*x*_*i*_ ∈ {1, 2, · · ·, 10)}. Intuitively, *m* denotes the proportion of migrants, and *σ* is the typical migration distance. Using this model, we examined the absorption time of a new mutation.

In Figure 3D–F, we plotted the absorption time of a new mutation that arises at subpopulation 1 (i.e., the left-most subpopulation) in a step environment. We assumed three different *m* while fixing *σ* = 5.0, allowing relatively long-distance migration. In all parameter sets examined, our theory accurately predicts the absorption time as long as *λ*_max_ < 0. We observed the loss of the typical trajectory (i.e., *λ*_max_ > 0) when *m* is small and both |*S*_1_| and |*S*_2_| are relatively large. Contrary to the two-population model, inapplicable cases arise also for cases with local adaptation. This is because when there are multiple demes where positive selection works on the focal allele, then multiple possible trajectories arise in a similar manner to Figure 1B.

We assumed different environmental distribution of the local pocket cases where *E*_2_ is located in the middle of the *E*_1_ (Figure S5). We also considered ten-island models where selection coefficient, subpopulation size, and migration rates are randomly drawn (Figure S6). Comparison between theory and the simulation results for these situations support the validity of our method even in complex multi-deme models.

### Effect of the pattern of environment on the stability of adaptive divergence

The stability of adaptive divergence depends not only on the strength of migration and selection, but also on the spatial distribution of environments. However, the effect of environmental distribution on the maintenance of variation remains elusive due to the simplified population structure in most previous analytical studies. In the fine-grained environments, locally adaptive alleles may go extinct locally for two reasons: (i) selection is averaged over adjacent environments and becomes weak, and (ii) random genetic drift is intensified owing to smaller local population size in each patchy environment. This instability at the local scale potentially increases the likelihood of loss of the locally adaptive alleles from the entire population. To quantify this effect, we apply our approximation method to two-dimensional space and examine how the stability of the adaptive divergence depends on the extent of environmental autocorrelation.

For this purpose, we designed two *in silico* experiments, using a space consisting of 8 × 8 subpopulations. In the first experiment, we compared three environmental landscapes of checker-board patterns with different size of component squares (Figure 4A). These landscapes have equal proportions of the two environments (*E*_1_ and *E*_2_) by area and symmetric selection coefficient (*s*_*i*_ = *S* in shaded patches while *s*_*i*_ = − *S* in white patches in Figure 4A), but differ in the spatial organization of the environments. Migration rate between demes is assumed to be

**Figure 4.**
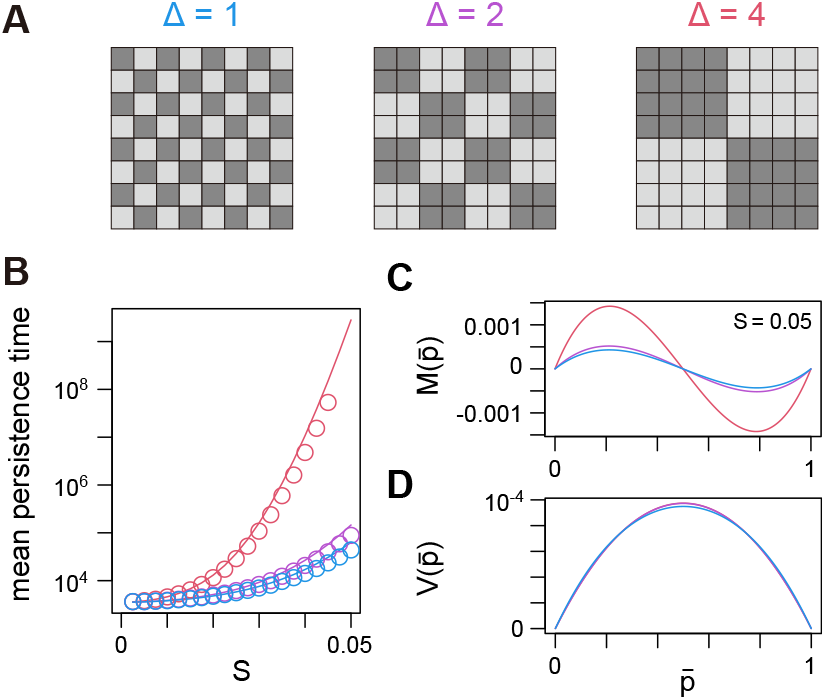
Effect of environmental patch size on the persistence of adaptive divergence. (A) Three checkerboard patterns examined. (B) Mean persistence time for each pattern. Lines are theoretical predictions while circles represent the simulation results (average over 100,000 replicates). (C, D) Values of 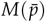 (C) and 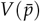 (D). Subpopulation size of each grid is 2*N*_*i*_ = 40, and *m* = 0.75, *σ* = 1.2 were assumed.

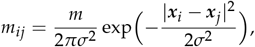

where ***x***_*i*_ is the coordinate of *i*th subpopulation. Using this model, we compare the persistence time of adaptive divergence starting from the coexistence of the two alleles at the stable deterministic equilibrium.

Figure 4B shows that, as expected, the persistence time is much longer in the more coarse-grained environments (i.e., large Δ). For example, the persistence time for *S* = 0.0275 in Δ = 4 is ≈ 53, 000, which is slightly longer than the time (≈ 44, 000) for *S* = 0.05 in Δ = 1. This pattern suggests that the spatial pattern of environment can have a similar effect as an 80% increase of selection strength on the maintenance of adaptive divergence. Theoretical predictions agree well with the simulation results for all cases.

To inspect whether the difference in stability arises from a change in selection or random genetic drift, we plotted the strength of the deterministic force 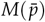 (Figure 4C) and the stochastic force 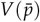 (Figure 4D) for the three cases. While all cases show a similar amount of stochasticity, the selection strength is greatly intensified in the coarse-grained environment, demonstrating that the intensified selection strength is the main cause of the increased stability in this scenario.

In the second experiment, we relax the assumption of an equal proportion of the two environments to examine the relative effect of the spatial pattern of environmental and total area of each environment. Let 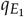 be the proportion of subpopulations in *E*_1_ where 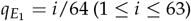. We then prepared the two types of landscape; a landscape where *E*_1_ environment is fully spatially clustered and 100 landscapes where environments are assigned at random across space (see the legend of Figure 5). For each landscape, we applied our theoretical method and obtained the expected persistence time of adaptive divergence starting from a deterministic equilibrium, if there is a stable polymorphic equilibrium. We repeated this procedure for all possible 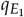. To remove any edge effects, the landscape is assumed to be toroidally wrapped such that the bottom (left) side is adjacent to the up (right) side of the landscape. Migration rate is given by 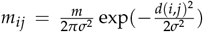 where *d*(*i, j*) denotes the shortest distance between *i*th and *j*th subpopulations in the wrapped landscape.

**Figure 5.**
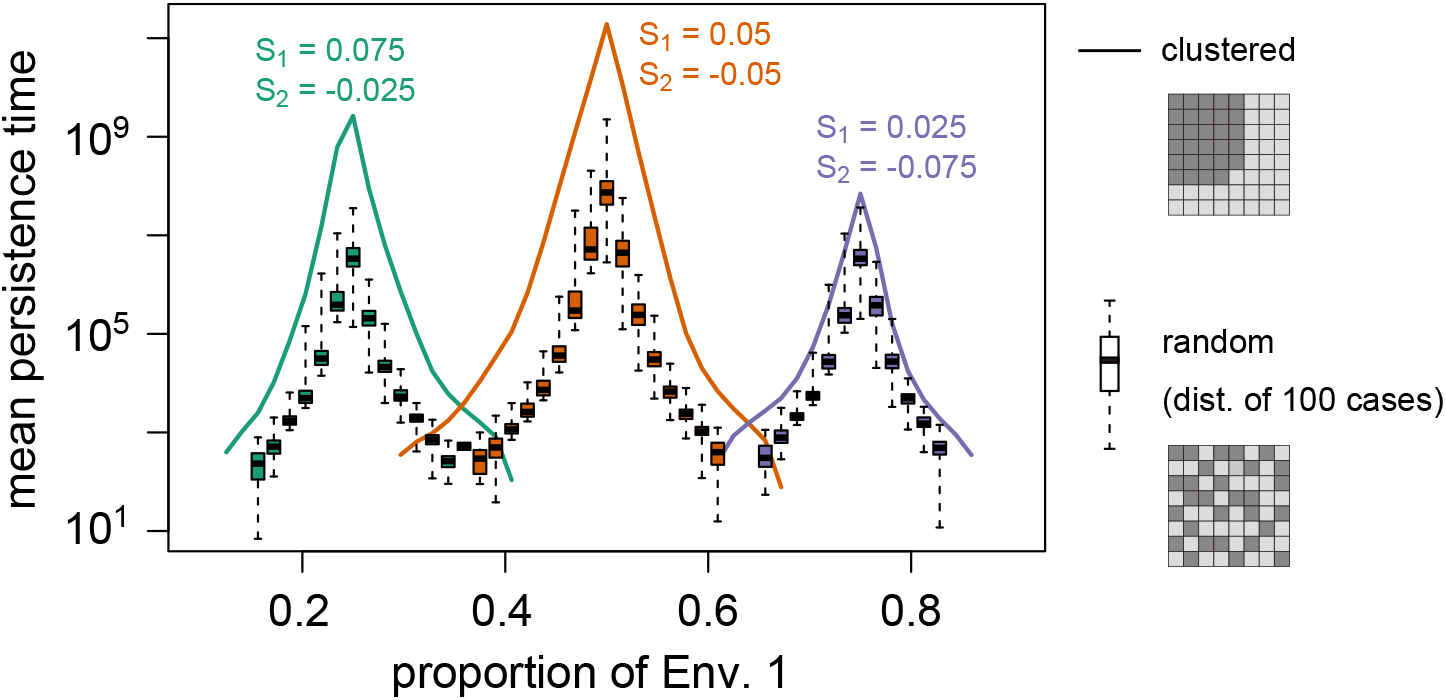
Effect of environmental distribution on the stability of adaptive divergence. Three selection coefficients were assumed: *S*_1_ = 0.075, *S*_2_ = − 0.025, *S*_1_ = 0.05, *S*_2_ = − 0.05, and *S*_1_ = 0.025, *S*_2_ = − 0.025. Solid lines show the results for the landscape in which *E*_1_ subpopulations are clustered, while box plots show the distribution of mean absorption time calculated for 100 randomly generated landscapes. Subpopulation size of each grid is 2*N*_*i*_ = 40, and *m* = 0.25, *σ* = 1.2 were assumed.

We compared the mean absorption time across various 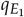 for three sets of selection coefficients (Figure 5). The range of 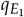 at which the polymorphism is maintained is determined by the relative strength of *S*_1_ and *S*_2_ such that 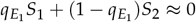. This pattern makes sense because average selection coefficient is zero for those conditions in the large migration limit (i.e., similar fitness for the two alleles). Since we assume relatively high migration rates where the stochasticity affects the polymorphism stability, the range of 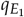 for persistent polymorphism is relatively narrow.

In general, the clustered environment enhances the stability of the polymorphic equilibrium. This effect is particularly large when *E*_1_ environment is not a common environment. When *S*_1_ = 0.075 and *S*_2_ = − 0.025, in which long-term coexistence is possible when 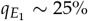, the difference in persistence time between the clustered and random landscapes roughly corresponds to a 3 ∼ 4% difference in 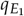, or a 15% relative increase/decrease of *E*_1_. When *S*_1_ = 0.05 and *S*_2_ = − 0.05, in which coexistence occurs in 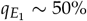, the difference between the two cases has the similar effect as a 4∼ 6% difference in 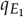, or 10% relative increase/decrease in *E*_1_. In contrast, when *S*_1_ = 0.025 and *S*_2_ = − 0.075, in which coexistence is possible when *E*_1_ is a common environment 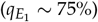, the difference in the two cases becomes relatively small (1 ∼ 3% difference of 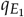). This pattern suggests that clustering for rare environments has a large effect on the maintenance of polymorphism, but a relatively weaker effect for common environments.

Although the above arguments are based on the theoretical prediction, we also ran simulations to check its performance. We randomly chosen 200 landscapes for each selection strength among ones where mean persistence time is predicted to be not so long (*T* < 10^7^), then compared the simulation and theory. Figure S7 shows that theory overall agrees with the simulation results. Although theory tends to overestimate the persistence time when adaptive divergence is stable, this difference is substantially less than an order of magnitude in all cases we studied, and does not affect the observed patterns in Figure 5. In summary, our theory is useful to see the overall patterns of the stability of adaptive divergence even when complex spatial heterogeneity is involved.

## Discussion

Our results show that, under sufficient migration, the core evolutionary processes can be summarized into a one-dimensional process and be analyzed in a similar manner as the one-population model. Importantly, this approximation can work even under complex spatial heterogeneity in selection strength, migration pattern, and subpopulation size, including some local adaptation cases. The present method brings two main benefits. First, we can understand the core evolutionary dynamics in an intuitive way because the one-dimensional process is easily visualized (see Figure 2C and Figure 4C, D). Second, this method provides a convenient way to calculate evolutionary dynamics for complex and realistic scenarios. Although the dynamics under complex settings can be examined by simulations in general, running many replicates becomes a computationally infeasible when we explore wide ranges of parameter space. This is particularly the case when local adaptation is involved because of the requirement for long computation times to assess persistence in scenarios with stable divergence. In contrast, our diffusion-based method can calculate the evolutionary quantities rapidly, which makes it possible to compare dynamics across a large number of different landscapes, as we showed in Figure 5.

By applying our method, we show that coarse-grained environments promote the maintenance of local adaptation. This is consistent with previous deterministic models showing that the polymorphic equilibrium becomes more stable as the width of environment increases (Slatkin 1973; Nagylaki 1975), while our method can quantify how this effect depends on various parameters accounting for stochasticity. In nature, organisms inhabit various types of environments where some selection pressures may have fine-grained patterns whereas others are relatively coarse grained. For example, in plant species, climate varies gradually over hundreds of kilometres across the habitat range, whereas soil type could vary over scales of kilometres or even meters. Our theory suggests that spatially varying selection favours the maintenance of polymorphism across a broader span of effect sizes for the more autocorrelated pressures (like climate) than the more fine-grained pressures (like soil condition). Consistent with this prediction, a recent report shows that climate has much larger contribution to local adaptation in *Arabidopsis thaliana* than the soil type (Ellis and Ågren 2024). We also observed that local adaptation for fine-grained environments is more vulnerable against change in environmental composition than that for coarse-grained environments, although this difference is at most modest in high migration (Figure 5). Therefore, some locally adaptive traits might be more sensitive to the environmental change like conversion of habitats.

These results also suggest the possibility to control the maintenance of adaptive genetic diversity by modifying the spatial arrangement of environments, even without changing the total area of each environment. A relevant situation would be the evolution of pest resistance in agriculture. Pesticide is widely used to improve agricultural yields, but its frequent use promotes the evolution of resistance that is locally advantageous under pesticide application (Kliot and Ghanim 2012; Hawkins *et al*. 2019). Our results suggest that such economically damaging local adaptation can be minimized if the cultivated/treated land is arranged in a fine-grained manner in the landscape, without reducing the total cultivation areas. Another relevant application might be maintenance of genetic diversity in urban areas: arranging parkland into a single large unit instead of many small ones could improve the maintenance of both urban- and rural-adapted ecotypes. Such an approach to design could increase the potential maintenance of diversity afforded by natural landscapes interspersed with urban ones.

Predicting the stability of adaptive divergence is important not only for managing genetic diversity, but also for interpreting genomic data undergoing local adaptation. In genomic analysis, peaks of genetic divergence between populations are widely used as a signal of local adaptation (Turner *et al*. 2005; Nosil 2012). The extent of divergence depends on the time since the establishment of locally adaptive alleles (Sakamoto and Innan 2019), thus divergence should be less pronounced when the coexistence of locally adaptive alleles is not so stable. Our theory predicts conditions under which the sustained local adaptation is probable so that we may observe detectable *F*_*ST*_ peak. Notably, the collapse of divergence is not taken into account in previous theories on *F*_*ST*_ peak in local adaptation as these studies assumed stable coexistence of selected alleles at the deterministic equilibrium after its establishment (Charlesworth *et al*. 1997; Yeaman *et al*. 2016; Sakamoto and Innan 2019). In future studies, these two frameworks may be integrated to predict the amount of genetic divergence accounting for the rare collapse of local adaptation, which would explain the stochastic variation of the observed genomic signal.

## Data availability

The codes for numerical calculation and simulation are available at https://github.com/TSakamoto-evo/multipopulation_approximation.

## Acknowledgments

We thank the members of Yeaman Lab. for helpful comments on the earlier version. This work was funded by JSPS KAKENHI Grant Number JP23KJ2158 to T.S; and grants from Alberta Innovates and NSERC Discovery to S.Y. The computing resource was provided by the Digital Research Alliance of Canada and Human Genome Center (the Univ. of Tokyo).

## Literature cited

Aeschbacher S, Bürger R. 2014. The effect of linkage on establishment and survival of locally beneficial mutations. Genetics. 197:317–336.

Akerman A, Bürger R. 2014. The consequences of gene flow for local adaptation and differentiation: a two-locus two-deme model. J. Math. Biol. 68:1135–1198.

Barton NH. 1987. The probability of establishment of an advantageous mutant in a subdivided population. Genet. Res. 50:35–40.

Barton NH, Whitlock MC. 1997. The evolution of metapopulations, In: Hanski I, Gilpin ME, editors, Metapopulation biology, Academic Press. San Diego. pp. 183–210.

Booker TR. 2024. The structure of the environment influences the patterns and genetics of local adaptation. Evolution Letters. 8:787–798.

Charlesworth B, Nordborg M, Charlesworth D. 1997. The effects of local selection, balanced polymorphism and background selection on equilibrium patterns of genetic diversity in subdivided populations. Genet. Res. 70:155–174.

Constable GW, McKane AJ. 2014. Fast-mode elimination in stochastic metapopulation models. Physical Review E. 89:032141.

Crow JF, Kimura M. 1970. An introduction to population genetics theory. Harper & Row.

Deakin MA. 1966. Sufficent conditions for genetic polymorphism. Am. Nat. 100:690–692.

Des Roches S, Pendleton LH, Shapiro B, Palkovacs EP. 2021. Conserving intraspecific variation for nature’s contributions to people. Nat. Ecol. Evol. 5:574–582.

Des Roches S, Post DM, Turley NE, Bailey JK, Hendry AP, Kinnison MT, Schweitzer JA, Palkovacs EP. 2018. The ecological importance of intraspecific variation. Nat. Ecol. Evol. 2:57–64.

Ellis TJ, Ågren J. 2024. Adaptation to soil type contributes little to local adaptation in an italian and a swedish ecotype of arabidopsis thaliana on contrasting soils. Biology Letters. 20:20240236.

Ewens WJ. 2004. Mathematical population genetics: theoretical introduction. volume 27. Springer.

Fisher RA. 1950. Gene frequencies in a cline determined by selection and diffusion. Biometrics. 6:353–361.

Haldane JBS. 1930. A mathematical theory of natural and artificial selection. (Part VI, Isolation.). Math. Proc. Camb. Philos. Soc. 26:220–230.

Haldane JBS. 1948. The theory of a cline. J. Genet. 48:277–284.

Hawkins NJ, Bass C, Dixon A, Neve P. 2019. The evolutionary origins of pesticide resistance. Biological Reviews. 94:135–155.

Kliot A, Ghanim M. 2012. Fitness costs associated with insecticide resistance. Pest Manag. Sci. 68:1431–1437.

Meek MH, Beever EA, Barbosa S, Fitzpatrick SW, Fletcher NK, Mittan-Moreau CS, Reid BN, Campbell-Staton SC, Green NF, Hellmann JJ. 2023. Understanding local adaptation to prepare populations for climate change. BioScience. 73:36–47.

Nagylaki T. 1975. Conditions for the existence of clines. Genetics. 80:595–615.

Nosil P. 2012. Ecological speciation. Oxford University Press.

Pollak E. 1966. On the survival of a gene in a subdivided population. J. Appl. Prob. 3:142–155.

Sakamoto T, Innan H. 2019. The evolutionary dynamics of a genetic barrier to gene flow: From the establishment to the emergence of a peak of divergence. Genetics. 212:1383–1398.

Slatkin M. 1973. Gene flow and selection in a cline. Genetics. 75:733–756.

Svoboda J, Tkadlec J, Kaveh K, Chatterjee K. 2023. Coexistence times in the moran process with environmental heterogeneity. Proc. R. Soc. A. 479:20220685.

Szép E, Sachdeva H, Barton NH. 2021. Polygenic local adaptation in metapopulations: A stochastic eco-evolutionary model. Evolution. 75:1030–1045.

Turner TL, Hahn MW, Nuzhdin SV. 2005. Genomic islands of speciation in anopheles gambiae. PLoS Biol. 3:e285.

Whitlock MC, Gomulkiewicz R. 2005. Probability of fixation in a heterogeneous environment. Genetics. 171:1407–1417.

Wright S. 1931. Evolution in mendelian populations. Genetics. 16:97.

Yeaman S, Aeschbacher S, Bürger R. 2016. The evolution of genomic islands by increased establishment probability of linked alleles. Mol. Ecol. 25:2542–2558.

Yeaman S, Otto SP. 2011. Establishment and maintenance of adaptive genetic divergence under migration, selection, and drift. Evolution. 65:2123–2129.

