## Supplementary information for "Maintenance of polymorphism in spatially heterogeneous environments"

### Supplementary materials for "Maintenance of polymorphism in spatially heterogeneous environments"

April 25, 2025

### 1 Construction of approximate diffusion process

#### 1.1 Overview of our approximation

This approximation presumes that migration is relatively strong and allele frequencies in different subpopulations are tightly coupled. Then, the evolutionary process is essentially a one-dimensional process, allowing us to apply a diffusion method. In the following, we first present intuitive ideas on (i) how to check whether this approximation is applicable, and (ii) how to constitute the one-dimensional diffusion process for applicable cases. Later, we propose a concrete method to calculate the diffusion coefficients.

The analysis takes the following four steps:

- (Step 1) Determine a one-dimensional path by deterministic differential equation:

$$\frac{dp_i}{dt} = s_i p_i(1-p_i) + \sum_{j\neq i} m_{ij}(p_j - p_i) \tag{S1}$$

  where initial conditions of $\bar{p} \ll 1$ and $1-\bar{p} \ll 1$ are applied (i.e., either of the two alleles is rare).

- (Step 2) Determine whether the evolutionary trajectory is convergent around the deterministic path. If so, starting from a frequency set close enough to the path, the evolution is expected to proceed along the path. Specifically, our aim is to distinguish convergent cases (Figure S1A) from divergent cases (Figure S1B). To check the convergence around a point on the deterministic path (point P), a plane that is orthogonal to the deterministic path ($l$) on which the point P resides is considered (Figure S1C). Starting from a point on the plane close to P (a point A in Figure S1C), the evolutionary direction ($\Delta\vec{p}$) is calculated by Equation S1 (vector $\overrightarrow{AB}$). This vector is projected on the plane (vector $\overrightarrow{AC}$), and the projected evolutionary system is constituted. We then apply a linear (perturbation) analysis around the point P to check the convergence. If the real part of leading eigenvalue of this linear system ($\lambda$) is negative, we consider that the evolutionary trajectory is convergent around the point P. Then, we check the convergence across the deterministic path and calculate $\lambda_{\max} = \max\lambda$ as an indicator of our method's applicability.

- (Step 3) For applicable cases, we calculate diffusion coefficients at each point on the deterministic path. Deterministic change ($\mathrm{E}[\Delta\bar{p}]$) is easily calculated from Equation S1. For stochastic change ($\mathrm{Var}[\Delta\bar{p}]$), we first evaluate how a small perturbation around the deterministic path affects the positional deviation along the deterministic path in a long run. In this derivation, we use an approximation based on linear dynamics around the deterministic path (see below for details).

- (Step 4) To expand the applicability to new mutations, we develop a method to determine the effective initial frequency $\bar{p}_0$ for new mutations with positive invasion fitness based on the branching process.

#### 1.2 Step 1

Calculate a deterministic path. As initial conditions, we assume either of the two alleles is rare. We assume that the initial frequency distribution reaches an equilibrium and follows the eigenvector corresponding to the leading eigenvector in the dynamic system of the rare allele.

Let us first consider the case with an allele A being rare ($p_i \sim \varepsilon \ll 1$). Ignore $\mathcal{O}(\varepsilon^2)$, Equation S1 becomes

$$\frac{dp_i}{dt} = s_i p_i + \sum_{j\neq i} m_{ij}(p_j - p_i), \tag{S2}$$

or in the matrix form,

$$\frac{d\vec{p}}{dt} = \boldsymbol{A}\vec{p}, \tag{S3}$$

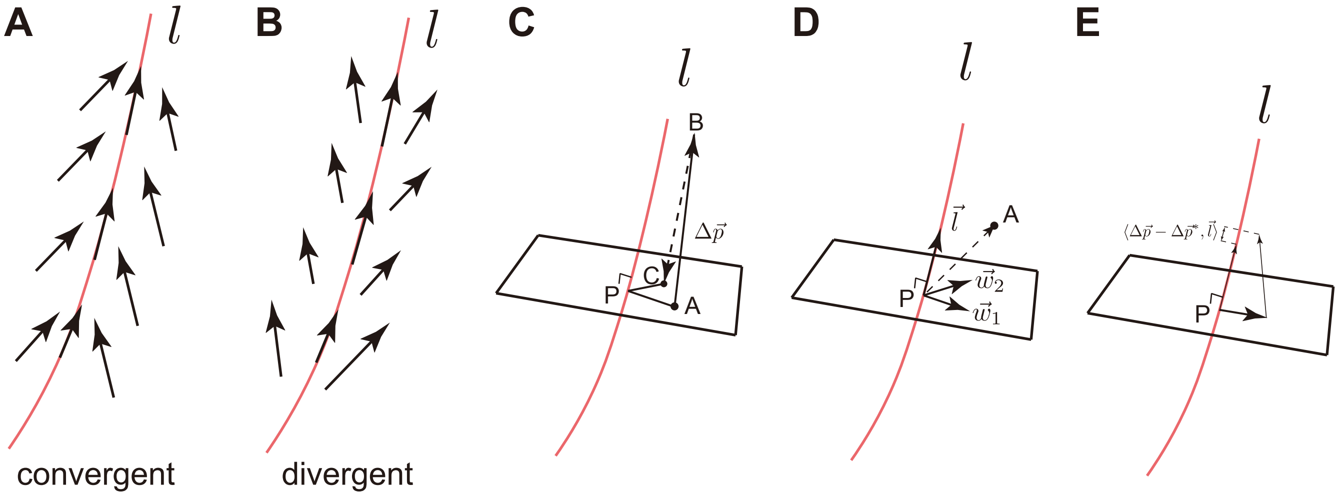


Figure S1: Illustration of the approximate diffusion construction.

where

$$a_{ij} = \begin{cases} m_{ij} & \text{for } i \neq j \\ s_i - \sum_{k \neq i} m_{ik} & \text{for } i = j. \end{cases}$$

If the leading eigenvalue of the matrix $\boldsymbol{A}$ is positive, allele A is invasive. For those cases, letting the corresponding eigenvector be $\vec{v}$, the initial frequency distribution is assumed to be $\vec{p} = \varepsilon \frac{\vec{v}}{|\vec{v}|}$ ($\varepsilon \ll 1$). Then, the deterministic path is calculated by Equation S1.

We applied the same method for the case with allele a being rare. If there is an internal stable equilibrium (i.e., both alleles are invasive), the two paths are connected at that equilibrium.

##### 1.3 Step 2

Let us consider a point P $(p_1^*, p_2^*, \cdots, p_n^*)$ on the deterministic path. Below, we use the symbol $*$ to emphasize that the focal point is on the deterministic path. A normalized vector along the deterministic path, $\vec{l}$, is calculated as

$$\vec{l} = \frac{1}{\sqrt{\sum_i (\Delta p_i^*)^2}} \begin{pmatrix} \Delta p_1^* \\ \vdots \\ \Delta p_n^* \end{pmatrix} \tag{S4}$$

where $\Delta p_i^* = s_i p_i^*(1 - p_i^*) + \sum_{j \neq i} m_{ij}(p_j^* - p_i^*)$.

We consider a plane that passes the point P and is orthogonal to $\vec{l}$ (Figure S1C). Letting a point A be on the plane close to the point P, its point coordinate is denoted by $p_i = p_i^* + \varepsilon_i$. The evolutionary direction from the point A ($\overrightarrow{AB}$ in Figure S1C) is calculated as

$$\begin{aligned} \frac{dp_i}{dt} &= \Delta p_i \\ &= s_i p_i (1 - p_i) + \sum_{j \neq i} m_{ij}(p_j - p_i). \end{aligned} \tag{S5}$$

We consider the projection of this dynamics onto the plane ($\overrightarrow{AC}$ in Figure S1C):

$$\begin{aligned} \frac{d\tilde{p}_i}{dt} &= \Delta p_i - \langle \Delta \vec{p}, \vec{l} \rangle l_i \\ &= \Delta p_i - \Big( \sum_j \Delta p_j l_j \Big) l_i. \end{aligned} \tag{S6}$$

Assuming $\varepsilon_i \ll 1$ and ignoring $\mathcal{O}(\varepsilon_i\varepsilon_j)$, we can write down the dynamics of $\varepsilon_i$ in the projected system:

$$\begin{aligned}\frac{d\varepsilon_i}{dt} &= s_i(1-2p_i^*)\varepsilon_i + \sum_{j\neq i} m_{ij}(\varepsilon_j - \varepsilon_i) - l_i \sum_j l_j \Big[s_j(1-2p_j^*)\varepsilon_j + \sum_{k\neq j} m_{jk}(\varepsilon_k - \varepsilon_j)\Big] \\ &= \Big[(1-l_i^2)\Big(s_i(1-2p_i^*) - \sum_{j\neq i} m_{ij}\Big) - \sum_{j\neq i} l_i l_j m_{ji}\Big]\varepsilon_i \\ &\quad + \sum_{j\neq i}\Big[(1-l_i^2)m_{ij} - l_i l_j s_j(1-2p_j^*) + l_i l_j \sum_{k\neq j} m_{jk} - \sum_{k\notin\{i,j\}} l_i l_k m_{kj}\Big]\varepsilon_j.\end{aligned} \tag{S7}$$

Let a matrix $\boldsymbol{X}$ be

$$x_{ij} = \begin{cases} (1-l_i^2)\Big(s_i(1-2p_i^*) - \sum_{j\neq i} m_{ij}\Big) - \sum_{j\neq i} l_i l_j m_{ji} & i = j \\ (1-l_i^2)m_{ij} - l_i l_j s_j(1-2p_j^*) + l_i l_j \sum_{k\neq j} m_{jk} - \sum_{k\notin\{i,j\}} l_i l_k m_{kj} & i \neq j. \end{cases} \tag{S8}$$

Using this matrix, the changes of $\varepsilon_i$ can be written in a matrix form as

$$\frac{d}{dt}\begin{pmatrix}\varepsilon_1\\ \vdots \\ \varepsilon_n\end{pmatrix} = \boldsymbol{X}\begin{pmatrix}\varepsilon_1\\ \vdots \\ \varepsilon_n\end{pmatrix}. \tag{S9}$$

In Equation S9, $(\varepsilon_1, \cdots, \varepsilon_n)^T$ is a coordinate vector with respect to the standard coordinate. Since we are now interested in the dynamics on the plane, we change a basis into $\vec{l}, \vec{\alpha}_1, \cdots, \vec{\alpha}_{n-1}$, where $(\vec{\alpha}_1, \cdots, \vec{\alpha}_{n-1})$ constitutes a normalized basis on the plane. Practically, we can find such a basis using the Gram-Schmidt process. Letting $\boldsymbol{U} = (\vec{l}, \vec{\alpha}_1, \cdots, \vec{\alpha}_{n-1})$, we can find a new coordinate vector $(b_0, \cdots, b_{n-1})^T$ with respect to the new basis:

$$\begin{pmatrix}b_0\\ \vdots \\ b_{n-1}\end{pmatrix} = \boldsymbol{U}^{-1}\begin{pmatrix}\varepsilon_1\\ \vdots \\ \varepsilon_n\end{pmatrix}.$$

Then, Equation S9 is rewritten as

$$\frac{d}{dt}\begin{pmatrix}b_0\\ \vdots \\ b_{n-1}\end{pmatrix} = \boldsymbol{Y}\begin{pmatrix}b_0\\ \vdots \\ b_{n-1}\end{pmatrix}, \tag{S10}$$

where $\boldsymbol{Y}$ is given by

$$\begin{aligned}\boldsymbol{Y} &= \boldsymbol{U}^{-1}\boldsymbol{X}\boldsymbol{U} \\ &= \begin{pmatrix} 0 & 0 & \cdots & 0 \\ y_{21} & & & \\ \vdots & & \tilde{\boldsymbol{Y}} & \\ y_{n1} & & & \end{pmatrix}.\end{aligned} \tag{S11}$$

The first row of $\boldsymbol{Y}$ should be 0 because we prohibit the movement along $\vec{l}$ in Equation S6. Here, the dynamics on the plane (i.e., $b_0 = 0$) is represented by

$$\frac{d}{dt}\begin{pmatrix}b_1\\ \vdots \\ b_{n-1}\end{pmatrix} = \tilde{\boldsymbol{Y}}\begin{pmatrix}b_1\\ \vdots \\ b_{n-1}\end{pmatrix}. \tag{S12}$$

Using the matrix $\tilde{\boldsymbol{Y}}$, we can determine the convergence of the evolutionary trajectories around the deterministic path. Let $\lambda_1, \cdots, \lambda_{n-1}$ be the eigenvalues of the matrix $\tilde{\boldsymbol{Y}}$. The condition for the convergence (i.e., $b_1, \cdots, b_{n-1} \to 0$ as $t \to \infty$) is presented as

$$\lambda \equiv \max \Re(\lambda_i) < 0. \tag{S13}$$

Let $\vec{\beta}_i$ be the right eigenvector of $\tilde{\boldsymbol{Y}}$ corresponding to the $i$th eigenvalue. We define a new basis on the plane, $\vec{w}_1, \cdots, \vec{w}_{n-1}$, by

$$(\vec{w}_1, \cdots, \vec{w}_{n-1}) = (\vec{\alpha}_1, \cdots, \vec{\alpha}_{n-1})(\vec{\beta}_1, \cdots, \vec{\beta}_{n-1}). \tag{S14}$$

Then, the coordinate vector with respect to the basis $\vec{w}_1, \cdots, \vec{w}_{n-1}$ is given by

$$\begin{pmatrix} c_1 \\ \vdots \\ c_{n-1} \end{pmatrix} = (\vec{\beta}_1, \cdots, \vec{\beta}_{n-1})^{-1} \begin{pmatrix} b_1 \\ \vdots \\ b_{n-1} \end{pmatrix}, \tag{S15}$$

and its dynamics are derived based on Equation S12:

$$\begin{aligned} \frac{d}{dt}\begin{pmatrix} c_1 \\ \vdots \\ c_{n-1} \end{pmatrix} &= (\vec{\beta}_1, \cdots, \vec{\beta}_{n-1})^{-1} \tilde{\boldsymbol{Y}} (\vec{\beta}_1, \cdots, \vec{\beta}_{n-1}) \begin{pmatrix} c_1 \\ \vdots \\ c_{n-1} \end{pmatrix} \\ &= \begin{pmatrix} \lambda_1 & 0 & \cdots & 0 \\ 0 & \lambda_2 & \cdots & 0 \\ \vdots & \vdots & \ddots & \vdots \\ 0 & 0 & \cdots & \lambda_{n-1} \end{pmatrix} \begin{pmatrix} c_1 \\ \vdots \\ c_{n-1} \end{pmatrix} \\ c_i(t) &= e^{\lambda_i t} c_i(0). \end{aligned} \tag{S16}$$

#### 1.4 Step 3

To constitute a diffusion process, we need to quantify the strength of deterministic and stochastic forces. Among these, the calculation of the deterministic term is straightforward. In the following, we specify positions on the deterministic path by average allele frequency $\bar{p}$:

$$\bar{p} = \frac{\sum_i N_i p_i}{\sum_i N_i}, \tag{S17}$$

where $N_i$ is a size of the $i$th subpopulation. Then, the deterministic change can be calculated as

$$\mathrm{E}[\Delta \bar{p}] = \frac{\sum_i \Delta p_i N_i}{\sum_i N_i}. \tag{S18}$$

The calculation of the stochastic term is more complicated. To calculate it, we need to know how the deviation from the deterministic path affects the directional change on the path after sufficient time. Below, we present an approximation method to quantify this effect.

Let a point A be near the point P that is on the deterministic path. Here, it is no longer assumed that the point A is on the orthogonal plane, but is assumed that it is not far from the plane (see Figure S1D). The vector $\overrightarrow{PA}$ represents a small perturbation, potentially due to the stochastic forces. The point coordinate of the point A is given by $p_i = p_i^* + \varepsilon_i$. Firstly, the vector $\overrightarrow{PA}$ is decomposed into

$$\begin{pmatrix} \varepsilon_1 \\ \vdots \\ \varepsilon_n \end{pmatrix} = c_0 \vec{l} + \sum_i c_i \vec{w}_i, \tag{S19}$$

where $\vec{w}_i$s are vectors defined in Equation S14.

In Equation S19, the long term effect from the first term is straightforward because this deviation shifts the position on the deterministic path by length $c_0$. For other terms, we need to consider the convergence process to the deterministic path. For this purpose, we reinterpret the analysis made in the step 2. In the step 2, we added the term of $-\langle \Delta\vec{p}, \vec{l}\rangle\vec{l}$ in the projection process to remove the evolutionary change along $\vec{l}$. Here, given that this subtraction should amount to $\langle \Delta\vec{p}^*, \vec{l}\rangle\vec{l}$ when the evolution starts from the point P, we can consider that $\langle \Delta\vec{p} - \Delta\vec{p}^*, \vec{l}\rangle\vec{l}$ as the extra subtraction arising from the deviation on the plane. In other words, we may consider $\langle \Delta\vec{p} - \Delta\vec{p}^*, \vec{l}\rangle\vec{l}$ as a shift in frequencies in the direction of $\vec{l}$ due to the deviation on the plane in the full system (i.e., the system without the projection) (Figure S1E). Thus, we can consider the summation of this term until the convergence as the directional change on the deterministic path. Noting that

$$\langle \Delta\vec{p}(t) - \Delta\vec{p}^*, \vec{l}\rangle = \sum_i l_i[s_i(1-2p_i^*)\varepsilon_i(t) + \sum_{j\neq i} m_{ij}(\varepsilon_j(t) - \varepsilon_i(t))] \tag{S20}$$

and the dynamics on the plane:

$$\begin{pmatrix} \varepsilon_1(t) \\ \vdots \\ \varepsilon_n(t) \end{pmatrix} = \sum_i c_i e^{\lambda_i t}\vec{w}_i, \tag{S21}$$

we can calculate the directional change in the direction of $\vec{l}$ due to the deviation on the plane as

$$\begin{aligned} \int_0^\infty \langle \Delta\vec{p}(t) - \Delta\vec{p}^*, \vec{l}\rangle dt &= \int_0^\infty \sum_i c_i e^{\lambda_i t}\xi_i dt \\ &= \sum_i -\frac{c_i\xi_i}{\lambda_i} \end{aligned} \tag{S22}$$

where $\xi_i = \sum_j l_j \left[s_j(1-2p_j^*)w_{ji} + \sum_{k\neq j} m_{jk}(w_{ki} - w_{ji})\right]$ and $w_{ji}$ is the $j$th component of $\vec{w}_i$. In the derivation, we used that $\Re(\lambda_i) < 0$, which should be satisfied for applicable cases.

Combining these two contributions, the amount of change along $\vec{l}$ due to the perturbation vector of $\overrightarrow{PA}$ can be approximated as

$$\delta\bar{p} = \frac{\sum_i l_i N_i}{\sum_i N_i}[c_0 - \sum_j \frac{c_j\xi_j}{\lambda_j}]. \tag{S23}$$

We are now be able to calculate the stochastic term of the diffusion process. The variance arising from the stochasticity in the subpopulation $i$ can be evaluated as

$$\mathrm{Var}_i[\Delta\bar{p}] = \left(\frac{\sum_j l_j N_j}{\sum_j N_j}[c_0^{(i)} - \sum_j \frac{c_j^{(i)}\xi_j}{\lambda_j}]\right)^2 \frac{p_i^*(1-p_i^*)}{2N_i} \tag{S24}$$

where

$$\begin{pmatrix} c_0^{(i)} \\ \vdots \\ c_{n-1}^{(i)} \end{pmatrix} = (\vec{l}, \vec{w}_1, \cdots, \vec{w}_{n-1})^{-1} \begin{pmatrix} \delta_{i1} \\ \vdots \\ \delta_{in} \end{pmatrix}. \tag{S25}$$

In this equation, $\delta_{ij} = 1$ for $i = j$, otherwise $\delta_{ij} = 0$. Summing up the stochasticity from all subpopulations, we can obtain:

$$\mathrm{Var}_{\mathrm{total}}[\Delta\bar{p}] = \sum_i \mathrm{Var}_i[\Delta\bar{p}]. \tag{S26}$$

We apply this method to each point on the deterministic path to determine $M(\bar{p}) = \mathrm{E}[\Delta\bar{p}]$ and $V(\bar{p}) = \mathrm{Var}_{\mathrm{total}}[\Delta\bar{p}]$. It should be noted that this method is not feasible for numerical calculation around internal stable equilibria because $|\vec{l}| \approx 0$ for those points. Practically, we can determine $M(\bar{p})$ and $V(\bar{p})$ by interpolation for those points.

##### 1.5 Step 4

In the above steps, we constructed an approximated diffusion process given that the initial position is very close to the deterministic path. However, this assumption can break down when we consider a new mutation which exists in only one of the subpopulations. To resolve this problem, we develop a simple method to determine the effective initial average frequency, $\bar{p}_0$, which is the allele frequency that shows the same long-term behavior as the focal case. We calculate $\bar{p}_0$ based on the establishment probability. This method is applicable to mutations with positive invasion fitness.

First, we consider the establishment probability in the approximate diffusion process. Let $M_0 = \lim_{\bar{p}\to 0} \frac{E[\Delta\bar{p}]}{\bar{p}}$ and $V_0 = \lim_{\bar{p}\to 0} \frac{\mathrm{Var}_{\mathrm{total}}[\Delta\bar{p}]}{\bar{p}}$. The establishment probability, which can be defined as the probability that a mutant escapes from immediate extinction, satisfies the following diffusion equation (Ewens, 2004):

$$\begin{aligned} 0 &= M_0\bar{p}\frac{du}{d\bar{p}} + \frac{1}{2}V_0\bar{p}\frac{d^2u}{d\bar{p}^2} \\ u(\bar{p}_0) &= 1 - \exp(-\frac{2M_0}{V_0}\bar{p}_0). \end{aligned} \tag{S27}$$

where $\bar{p}_0$ is the effective initial allele frequency.

Another method to calculate the establishment probability is to use the multi-type branching process (Pollak, 1966; Barton, 1987):

$$1 - P_i = \exp\Big(-\exp(s_i)\Big[\big(1 - \sum_{j\neq i} m_{ij}\big)P_i + \sum_{j\neq i}\frac{N_j m_{ji}}{N_i}P_j\Big]\Big) \tag{S28}$$

where $P_i$ is the establishment probability of an allele in the $i$th subpopulation. This probability can be determined by numerical iteration.

By equalizing these two probabilities, we can determine the effective initial frequency when the evolutionary process starts from a mutation in the $i$th population as

$$\begin{aligned} u(\bar{p}_0) &= P_i \\ \bar{p}_0 &= -\frac{V_0}{2M_0}\log(1 - P_i). \end{aligned} \tag{S29}$$

#### 2 Calculation of mean sojourn time and mean absorption time

Using the derived diffusion coefficients, mean sojourn time and mean absorption time were derived by diffusion methods (Ewens, 2004).

Letting $p_0$ be the initial frequency, the mean sojourn time density at frequency $p$ is calculated as

$$t(p;p_0) = \begin{cases} \frac{2}{V(p)G(p)}\frac{\int_{p_0}^1 G(q)dq}{\int_0^1 G(q)dq}\int_0^p G(q)dq & 0 \le p \le p_0 \\ \\ \frac{2}{V(p)G(p)}\frac{\int_0^{p_0} G(q)dq}{\int_0^1 G(q)dq}\int_p^1 G(q)dq & p_0 \le p \le 1, \end{cases} \tag{S30}$$

where $G(p) = \exp(-\int \frac{2M(p)}{V(p)}dp)$. For new mutations, we can further assume that $p_0 \ll 1$, leading to

$$t(p;p_0) \approx \frac{2p_0}{V(p)G(p)}\frac{\int_p^1 G(q)dq}{\int_0^1 G(q)dq}. \tag{S31}$$

We substitute $M(\bar{p})$, $V(\bar{p})$, and $\bar{p}_0$ into these equations to obtain mean sojourn time.

Mean absorption time is calculated from mean sojourn time:

$$\begin{aligned} T &= \int_{\varepsilon}^{1-\varepsilon} t(p; p_0) dp \\ &\approx \sum_{i=1}^{L-1} t\Big(\frac{i}{L}; p_0\Big) \frac{1}{L} \end{aligned} \tag{S32}$$

where $L$ is the number of discretization points in the numerical calculation. In this paper, $L = 1{,}000$ was used for all numerical results.

### 3 Performance of previous methods

As far as we know, Whitlock and Gomulkiewicz (2005) and Constable and McKane (2014) so far attempted to approximate the dynamics of multi-population model by one-dimensional diffusion. Although the construction procedure of these two theories seems different, they share the basic strategy. First, assuming the difference in allele frequencies among subpopulations is small, the evolutionary dynamics are assumed to be mainly determined by the average selection coefficient. Second, the effect of small allele frequency divergence is incorporated to adjust the strength of selection. Because of this similarity, the constructed diffusion process is very similar between the two methods.

To illustrate this point, we focus on the two population model with symmetric migration as an example. Let $s_i$ be the selection coefficient in subpopulation $i$. We assume that the migration rate and the population sizes are the same for two subpopulations ($m_{12} = m_{21} = m$ and $N_1 = N_2 = N$). Using Equations 16 and 17 in Whitlock and Gomulkiewicz (2005), $M(\bar{p}) = \mathrm{E}[\Delta\bar{p}]$ and $V(\bar{p}) = \mathrm{Var}[\Delta\bar{p}]$ are calculated as

$$\begin{aligned} M_{WG}(\bar{p}) &\approx \frac{s_1+s_2}{2}(1-F_{ST})\bar{p}(1-\bar{p}) + \frac{(s_1-s_2)^2}{8m}\bar{p}(1-\bar{p})(1-2\bar{p}) \\ V_{WG}(\bar{p}) &\approx \frac{\bar{p}(1-\bar{p})}{4N}(1-F_{ST}). \end{aligned} \tag{S33}$$

where $F_{ST} \approx \frac{1}{1+8Nm}$. Similarly, using Equations 23 and 33 in Constable and McKane (2014), diffusion coefficients are calculated as

$$\begin{aligned} M_{CM}(\bar{p}) &= \frac{s_1+s_2}{2}\bar{p}(1-\bar{p}) + \frac{(s_1-s_2)^2}{8m}\bar{p}(1-\bar{p})(1-2\bar{p}) \\ V_{CM}(\bar{p}) &\approx \frac{\bar{p}(1-\bar{p})}{4N}. \end{aligned} \tag{S34}$$

As clear from Equations S33 and S34, the resulting diffusion equations are almost identical. The only difference is the incorporation of $(1 - F_{ST})$ in Whitlock and Gomulkiewicz (2005). Since these methods assumed relatively strong migration, $F_{ST}$ should be $\ll 1$ so that the two equations give essentially the same results. It should be noted that while the method of Whitlock and Gomulkiewicz (2005) is applicable to island models with two environments, the method of Constable and McKane (2014) is more general allowing arbitrary choice of $s_i$, $m_{ij}$, and $N_i$.

To check the performance, we plotted the absorption time predicted by each method (Figure S2). For comparison with our method, please see Figure 3A-C in the main text. Figure S2 shows that the two methods have similar qualitative patterns: performance is good as long as $s_1 \approx s_2$, but it becomes poor when the two selection coefficients are moderately different and migration is infrequent. Notably, these methods cannot be applied to cases with divergent selection, as expected from their assumptions that frequency divergence is small and the evolutionary process is still close to the neutral one. Given the good performance of our method in wide range of parameters (Figure 3A-C), we argue that our method is useful in more general situations including the local adaptation case.

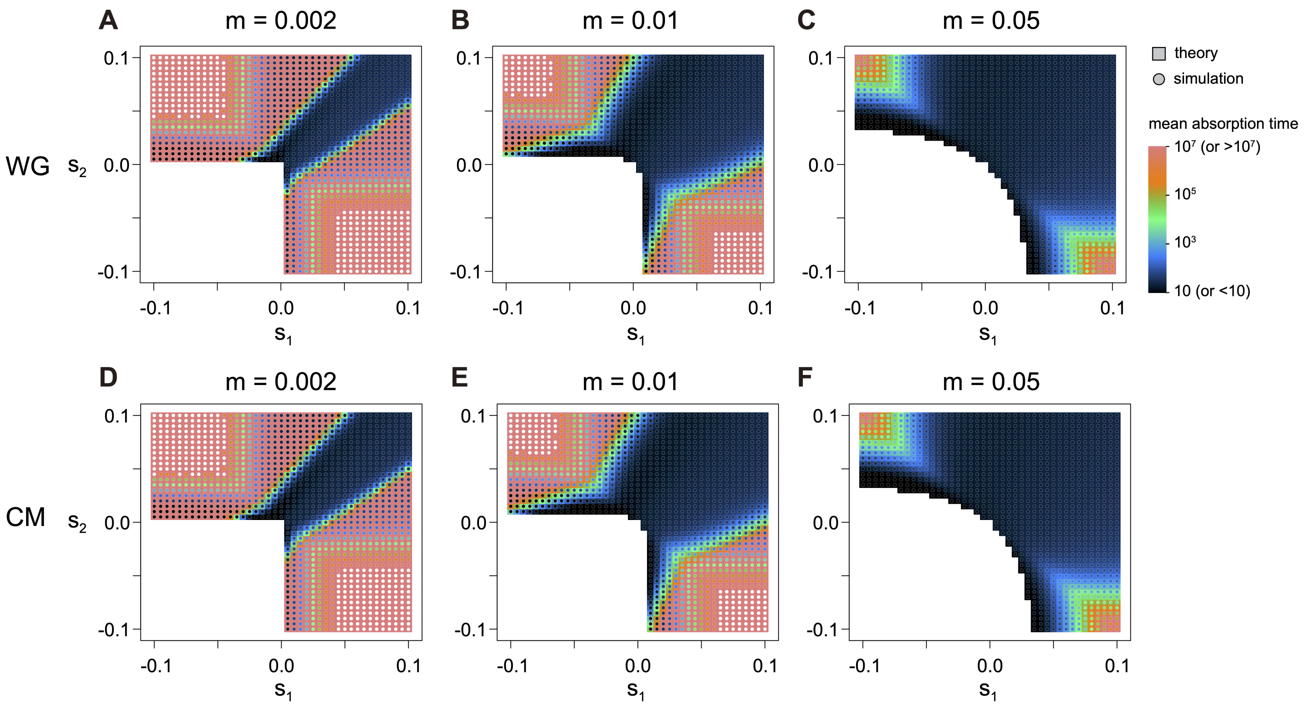


Figure S2: Performance of the previous methods in the two-population model. The same parameter range as Figure 3A-C was assumed. Theoretical prediction by Whitlock and Gomulkiewicz (2005) is plotted in panels A-C, while prediction by Constable and McKane (2014) is plotted in panels D-F.

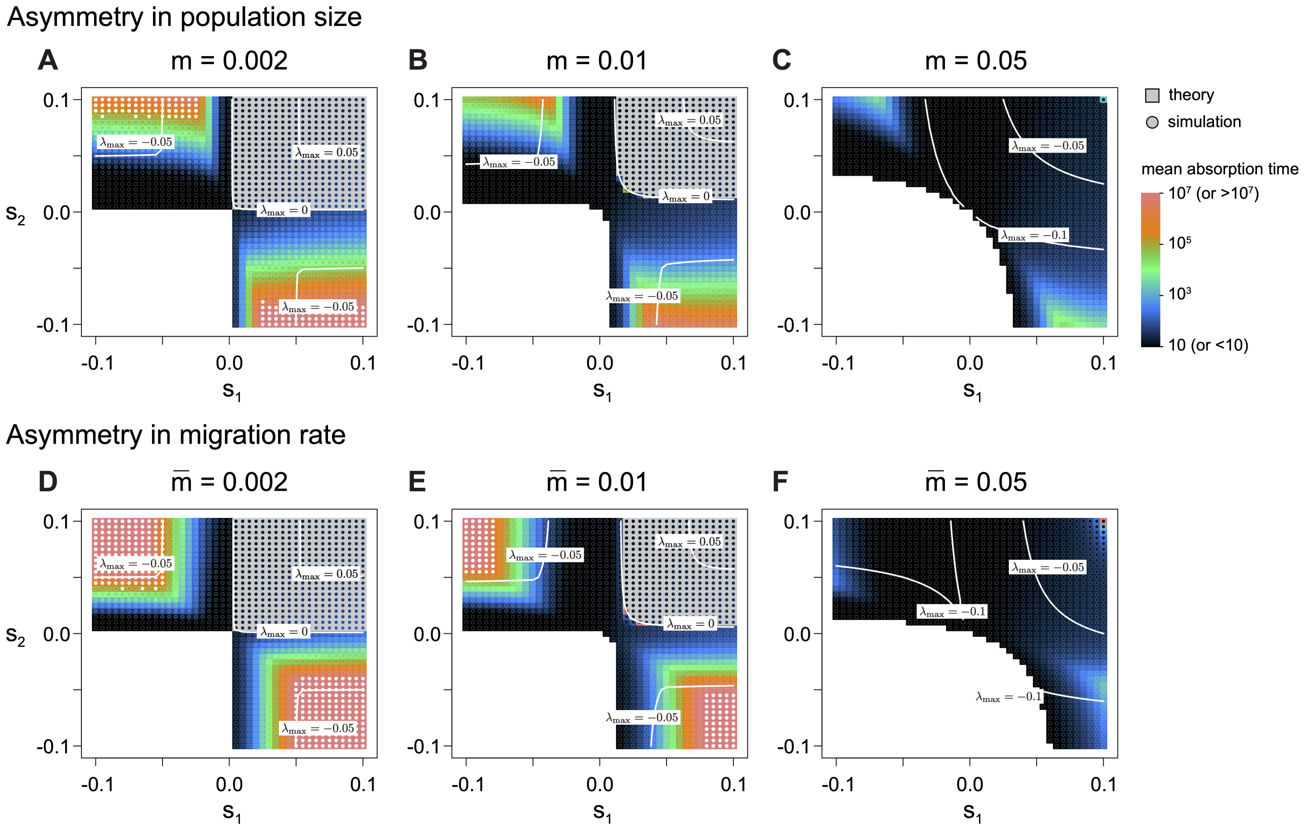


Figure S3: Effect of asymmetry in population size and migration in the two-population model. In panels (A-C), asymmetric population sizes ($2N_1 = 500, 2N_2 = 100$) are assumed. In panels (D–F), asymmetric migration rates ($m_{12} = 3m_{21}$) are assumed with $2N_i = 200$.

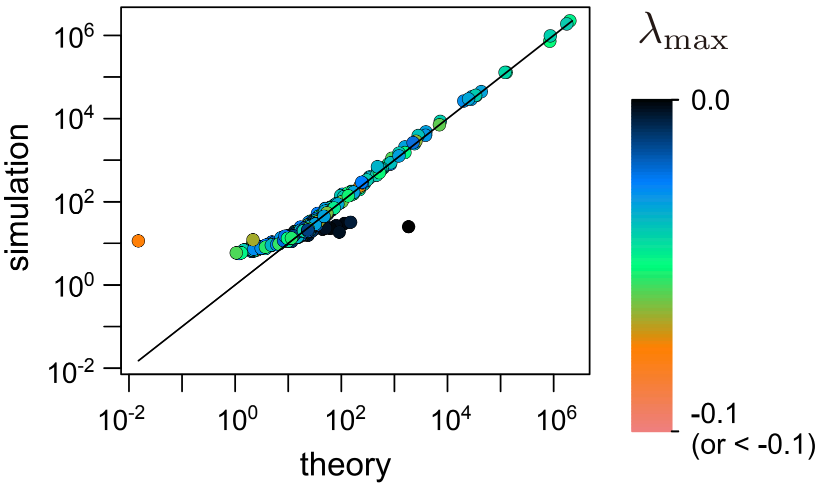


Figure S4: Mean absorption time in the two-population model with randomly assigned parameters. $s_1, s_2 \sim \text{Uniform}(-0.05, 0.05)$, $N_1, N_2 \sim \text{Uniform}(50, 250)$, and $\log(m_{12}), \log(m_{21}) \sim \text{Uniform}(\log(0.005), \log(0.05))$. Only cases with theoretical applicability (i.e., positive invasion fitness of a mutant allele and $\lambda_{\max} < 0$) are shown.

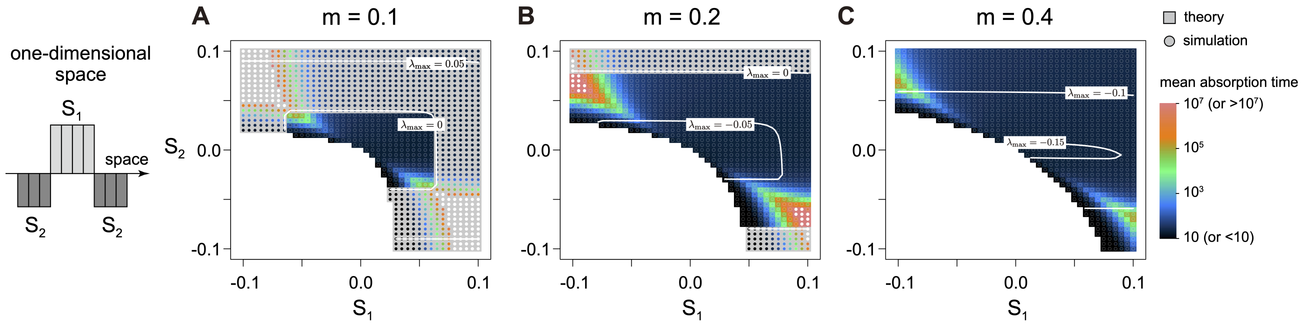


Figure S5: Mean absorption time in the local pocket environment. Four subpopulations in the middle have $E_1$, while the other subpopulation have $E_2$. $2N_i = 100$ and $\sigma = 5.0$ were assumed.

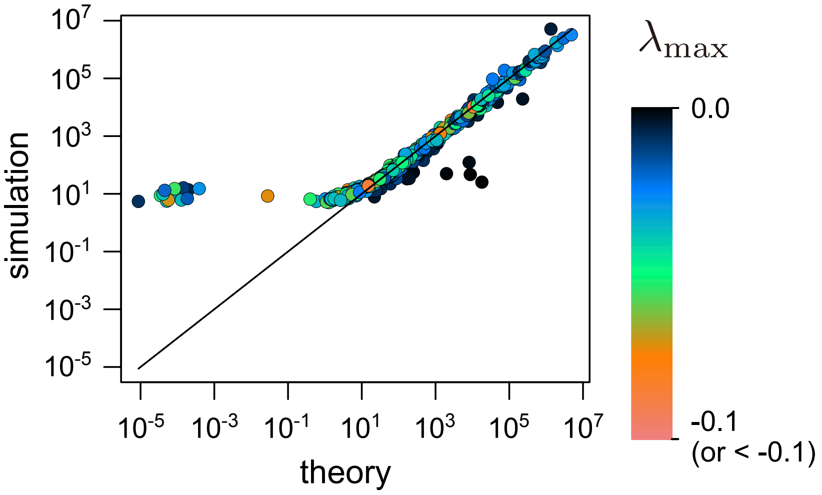


Figure S6: Mean absorption time in the ten-island model with randomly assigned parameters. $s_i \sim \text{Uniform}(-0.1, 0.1)$, $N_i \sim \text{Uniform}(25, 100)$, and $\log(m_{ij}) \sim \text{Uniform}(\log(0.002), \log(0.05))$. Only cases with theoretical applicability (i.e., positive invasion fitness of a mutant allele and $\lambda_{\max} < 0$) are shown.

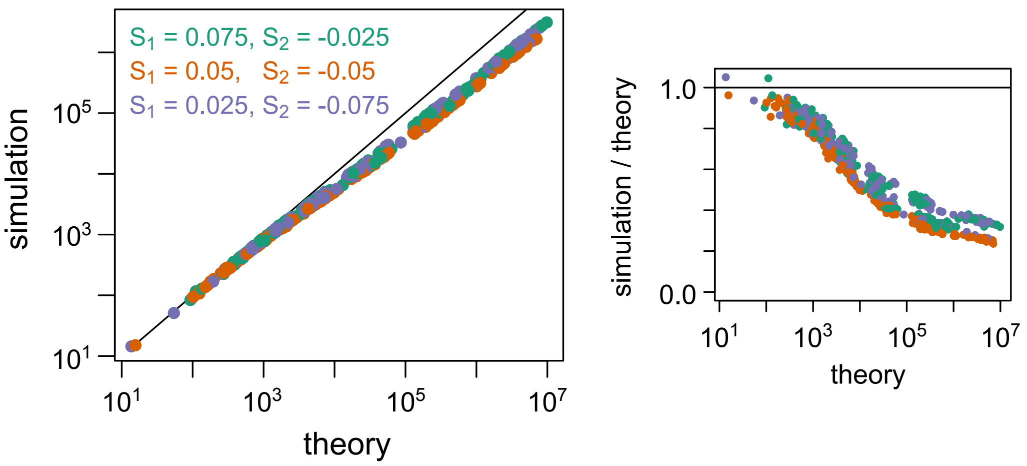


Figure S7: Mean absorption time in 8×8 landscapes. Landscapes are randomly chosen from ones that were used in Figure 5 in the main text. For each selection strength, 200 environmental distributions are selected among cases where the persistence time in theory is shorter than $10^7$ generations. For each case, 1,000 simulations were run to obtain a mean value.

### References

Barton NH. 1987. The probability of establishment of an advantageous mutant in a subdivided population. Genet. Res.. 50:35–40.

Constable GW, McKane AJ. 2014. Fast-mode elimination in stochastic metapopulation models. Phys. Rev. E. 89:032141.

Ewens WJ. 2004. *Mathematical population genetics: theoretical introduction*. volume 27. Springer.

Pollak E. 1966. On the survival of a gene in a subdivided population. J. Appl. Prob.. 3:142–155.

Whitlock MC, Gomulkiewicz R. 2005. Probability of fixation in a heterogeneous environment. Genetics. 171:1407–1417.
